# Age and life expectancy clocks based on machine learning analysis of mouse frailty

**DOI:** 10.1101/2019.12.20.884452

**Authors:** Michael B Schultz, Alice E Kane, Sarah J Mitchell, Michael R MacArthur, Elisa Warner, James R. Mitchell, Susan E Howlett, Michael S Bonkowski, David A Sinclair

**Author notes:** These authors contributed equally (Michael B Schultz, Alice E Kane). Corresponding author: David A. Sinclair.

## Abstract

The identification of genes and interventions that slow or reverse aging is hampered by the lack of non-invasive metrics that can predict life expectancy of pre-clinical models. Frailty Indices (FIs) in mice are composite measures of health that are cost-effective and non-invasive, but whether they can accurately predict health and lifespan is not known. Here, mouse FIs were scored longitudinally until death and machine learning was employed to develop two clocks. A random forest regression was trained on FI components for chronological age to generate the FRIGHT (<u>Fr</u>ailty Inferred <u>G</u>eriatric <u>H</u>ealth <u>T</u>imeline) clock, a strong predictor of chronological age. A second model was trained on remaining lifespan to generate the AFRAID (<u>A</u>nalysis of <u>Frai</u>lty and <u>D</u>eath) clock, which accurately predicts life expectancy and the efficacy of a lifespan-extending intervention up to a year in advance. Adoption of these clocks should accelerate the identification of novel longevity genes and aging interventions.

## INTRODUCTION

Aging is a biological process that causes physical and physiological deficits over time, culminating in organ failure and death. For species that experience aging, which includes nearly all animals, its presentation is not uniform; individuals age at different rates and in different ways. *Biological age* is an increasingly utilized concept that aims to more accurately reflect aging in an individual than the conventional *chronological age*. Biological measures that accurately predict health and longevity would greatly expedite studies aimed at identifying novel genetic and pharmacological disease and aging interventions.

Any useful biometric or biomarker for biological age should track with chronological age and should serve as a better predictor of remaining longevity and other age-associated outcomes than does chronological age alone, even at an age when most of a population is still alive. In addition, its measurement should be non-invasive to allow for repeated measurements without altering the health or lifespan of the animal measured (Butler et al. 2004). In humans, biometrics and biomarkers that meet at least some of these requirements include physiological measurements such as grip strength or gait (Rantanen et al. 2000; Bittner et al. 1993), measures of the immune system (Alpert et al. 2019; Martínez de Toda et al. 2019), telomere length (Mather et al. 2011), advanced glycosylation end-products (Krištić et al. 2014), levels of cellular senescence (Wang and Dreesen 2018), and DNA methylation clocks (Horvath 2013b). DNA methylation clocks have been adapted for mice but unfortunately these clocks are currently expensive, time consuming, and require the extraction of blood or tissue.

Frailty index assessments in humans are strong predictors of mortality and morbidity, outperforming other measures of biological age including DNA methylation clocks (Kim et al. 2017; Horvath and Raj 2018). Frailty indices quantify the accumulation of up to 70 health-related deficits, including laboratory test results, symptoms, diseases and standard measures such as activities of daily living (Searle et al. 2008; Mitnitski, Mogilner, and Rockwood 2001). The number of deficits an individual shows is divided by the number of items measured to give a number between 0 and 1, in which a higher number indicates a greater degree of frailty. The frailty index has been recently reverse-translated into an assessment tool for mice which includes 31 non-invasive items across a range of systems (Whitehead et al. 2014). The mouse frailty index is strongly associated with chronological age (Whitehead et al. 2014; Kane et al. 2018), correlated with mortality and other age-related outcomes (Feridooni et al. 2017; Rockwood et al. 2017), and is sensitive to lifespan-altering interventions (Kane et al. 2016). However, the power of the mouse frailty index to model biological age or predict life expectancy for an individual animal has not yet been explored.

In this study, we tracked frailty longitudinally in a cohort of aging male mice from 21 months of age until their natural deaths and employed machine learning algorithms to build two clocks: FRIGHT age, designed to model chronological age, and the AFRAID clock, which was modelled to predict life expectancy. FRIGHT age reflects apparent chronological age better than FI alone, while the AFRAID clock predicts life expectancy at multiple ages. These clocks were then tested for their predicitve power on cohorts of mice treated with interventions known to extend healthspan or lifespan, enalapril and methionine restriction. They accurately predicted increased healthspan and lifespan, demonstrating that an assessment of non-invasive biometrics in interventional studies can greatly accelerate the pace of discovery.

## RESULTS

### Frailty correlates with and is predictive of age

We measured FI scores (Figure S1) approximately every six weeks in a population of naturally aging male C57BL/6Nia mice (n=51) until the end of their lives. These mice had a normal lifespan, with a median survival of 30 months and a maximum (90^th^ percentile) of 36 months (Figure 1a, Figure S2). As expected, FI scores increased with age from 21 to 36 months at the population level (Figure 1b). At the individual level, frailty trajectories displayed significant variance, representative of the variability in how individuals experience aging within a population of inbred animals (Figure 1c). As FI score was well-correlated with chronological age, we sought to determine the degree to which FI score could model chronological and biological age. We performed a linear regression on FI score for age with a training dataset and evaluated its accuracy on a testing dataset (Figure 1d-e). FI score was able to predict chronological age with a median error of 1.8 months and an r-squared value of 0.642 (p=7.3e^-20^). We hypothesized that the error may be representative of biological age, with healthier individuals having a predicted age younger than their true age. We calculated this difference between predicted age and true age, termed delta age, and used remaining time until death as our primary outcome to compare with. For some individual age groups (24, 34.5 and 36 months), delta age did indeed have a negative correlation with survival, with “younger” mice (those with a negative delta age) living longer at each individual age than “older” mice (those with a positive delta age) (Figure 1f, Table 1). For other groups this correlation is a trend, and more power may detect an association (Table 1). This suggests that the FI score is able to detect variation in predicted chronological age for mice of the same actual age, and this may represent biological age.

**Table 1.** Correlation coefficients (r^2^) and p values for correlation between survival and delta age determined by either FI score or FRIGHT age, at individual ages.

| Age | n | Delta Age –<br>FI Score<br>(Figure 1f) |  | Delta Age -<br>FRIGHT Age<br>(Figure 3g) |  | Survival –<br>AFRAID Clock<br>(Figure 4g) |  |
| --- | --- | --- | --- | --- | --- | --- | --- |
| | | $r^2$ | p value | $r^2$ | p value | $r^2$ | p value |
| 21.0 | 24 | 0.021 | 0.623 | 0.010 | 0.728 | 0.035 | 0.519 |
| 22.5 | 27 | 0.039 | 0.537 | 0.005 | 0.831 | 0.015 | 0.708 |
| 24.0 | 20 | 0.390 | 0.023* | 0.296 | 0.054 | 0.447 | 0.012* |
| 25.5 | 29 | 0.142 | 0.124 | 0.012 | 0.659 | 0.121 | 0.157 |
| 27.0 | 43 | 0.103 | 0.109 | 0.004 | 0.753 | 0.218 | 0.016* |
| 28.5 | 36 | 0.109 | 0.124 | 0.031 | 0.422 | 0.191 | 0.037* |
| 30.0 | 32 | 0.062 | 0.291 | 0.019 | 0.567 | 0.363 | 0.004* |
| 31.5 | 25 | 0.003 | 0.859 | 0.011 | 0.706 | 0.229 | 0.071 |
| 33.0 | 18 | 0.305 | 0.063 | 0.276 | 0.079 | 0.292 | 0.069 |
| 34.5 | 11 | 0.686 | 0.021* | 0.661 | 0.026* | 0.653 | 0.028* |
| 36.0 | 6 | 0.881 | 0.018* | 0.396 | 0.256 | 0.201 | 0.448 |

**Figure 1.**
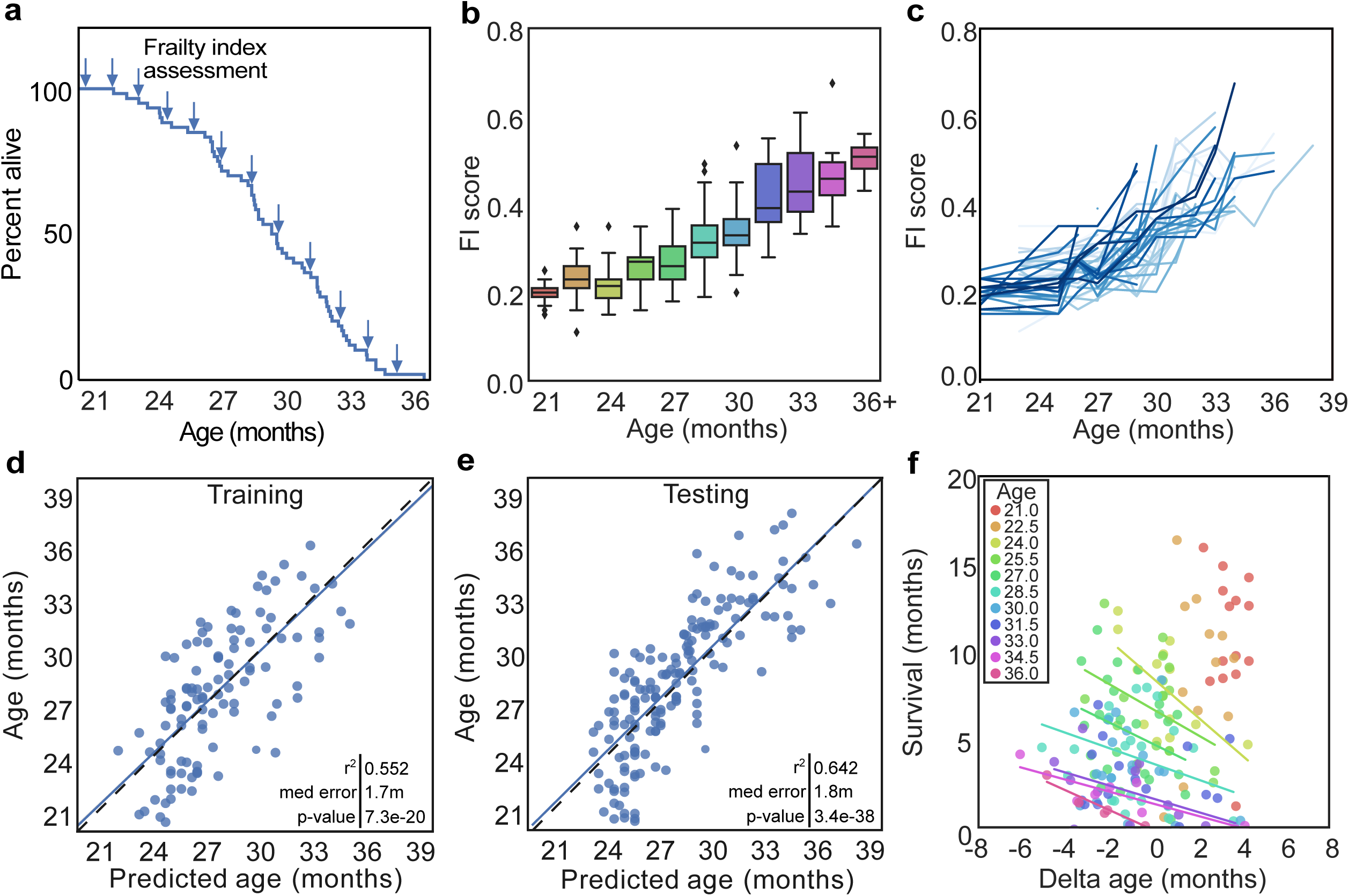
Frailty correlates with and is predictive of age in mice. (a) Kaplan-Meier survival curve for male C57BL/6 mice (n=51) assessed longitudinally for frailty index (indicated by arrows). (b) Box and whisker plots for mean frailty index (FI) score for mice from 21 to 36 months of age. (c) FI score trajectories for each individual mouse from 21 months until death. (d) Univariate regression of FI score for chronological age on a training dataset, and (e) a testing dataset. (f) Residuals of the regression (delta age), plotted against survival for individual ages. Regression lines are only graphed for ages where there is an r^2^ value > 0.1.

### Individual frailty items vary in their correlation with age

While a simple linear regression on overall frailty score was somewhat predictive of age, we hypothesized that by differentially weighting individual metrics, we could build a more predictive model, as has been done with various CpG sites to build methylation clocks (Horvath 2013b). To this end, we calculated the correlation between each individual frailty index item and chronological age (Table 2). Some parameters, such as tail stiffening, breathing rate/depth, gait disorders, hearing loss, kyphosis, and tremor, are strongly correlated (r^2^ > 0.35, p < 1e^-30^) with age (Figure 2), while others show very weak or no correlation with age (Table 2, Supplementary Figure 2). The fact that some parameters were very well correlated and others poorly correlated suggested that by weighting items we could build an improved model for biological age prediction.

**Table 2.** Correlation coefficients (r^2^) and p values for individual frailty items with chronological age.

| **Item** | **$r^2$** | **p value** |
| --- | --- | --- |
| FI Score | 0.61 | <0.001 |
| Tail stiffening | 0.58 | <0.001 |
| Breathing rate/depth | 0.50 | <0.001 |
| Gait disorders | 0.42 | <0.001 |
| Hearing loss | 0.38 | <0.001 |
| Kyphosis | 0.38 | <0.001 |
| Tremor | 0.38 | <0.001 |
| Body condition score | 0.26 | <0.001 |
| Forelimb grip strength | 0.12 | <0.001 |
| % twc | 0.12 | <0.001 |
| Menace reflex | 0.11 | <0.001 |
| Alopecia | 0.10 | <0.001 |
| Tumors | 0.08 | <0.001 |
| Diarrhea | 0.05 | <0.001 |
| Penile prolapse | 0.05 | <0.001 |
| Microphthalmia | 0.05 | <0.001 |
| Dermatitis | 0.05 | <0.001 |
| Rectal prolapse | 0.04 | <0.001 |
| Distended abdomen | 0.04 | <0.001 |
| Eye discharge/swelling | 0.04 | <0.001 |
| Coat condition | 0.04 | <0.001 |
| Body weight score | 0.02 | 0.01 |
| Threshold % rwc | 0.01 | 0.05 |
| Loss of fur color | 0.01 | 0.06 |
| Piloerection | 0.01 | 0.07 |
| Mouse grimace scale | 0.01 | 0.08 |
| Vestibular disturbance | 0.01 | 0.16 |
| Vision loss | 0.00 | 0.30 |
| Loss of whiskers | 0.00 | 0.33 |
| % rwc | 0.00 | 0.49 |
| Cataracts | 0.00 | 0.68 |
| Corneal capacity | 0.00 | 0.72 |
| Nasal discharge | 0.00 | 1.00 |

Twc, total weight change; rwc, recent weight change.

**Figure 2.**
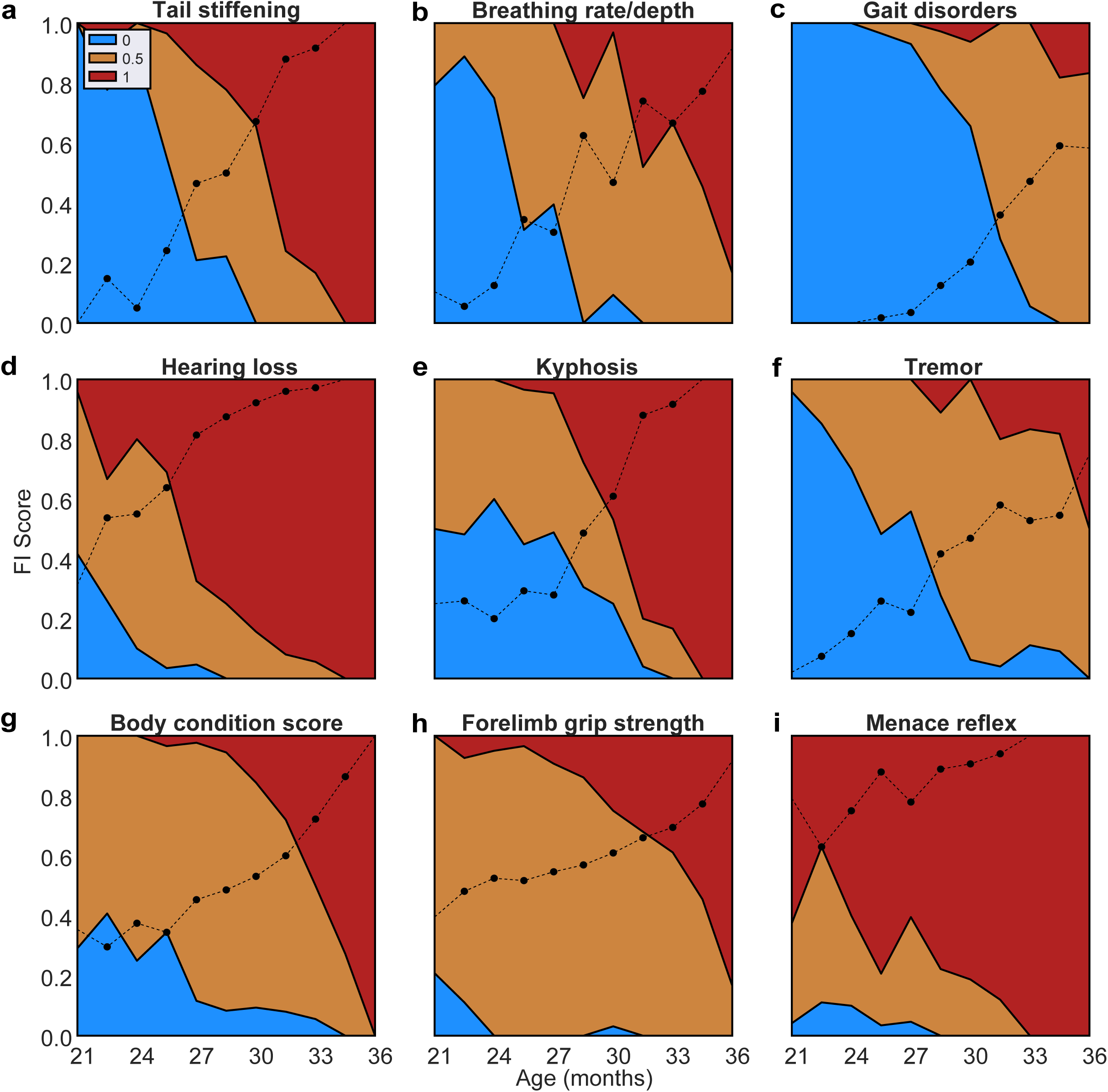
Individual FI items vary in their correlation with age. Mean scores across all mice (black line) for the top nine individual items of the frailty index that were correlated with chronological age. Colors indicate proportion of mice at each age with each score (0, blue; 0.5, orange, 1, red).

### Multivariate regressions of individual frailty items to predict age (FRIGHT age)

We compared FI score as a single variable and four types of multivariate linear regression models to predict chronological age: simple least squares regression, elastic net regression, random forest regression and the Klemera-Doubal biological age estimation method (Klemera and Doubal 2006). We employed the bootstrap method on the training dataset to compare models. Only frailty items that had a significant, even if weak, correlation with age (p<0.05) were included in the analysis (see Table 2). The multivariate models, particularly elastic net, the Klemera-Doubal method and random forest were superior to FI as a single variable, with lower median error (p<0.0001, F=50.46, df=495), higher r^2^ values (p<0.0001, F=74.38, df=496), and smaller p-values (p<0.0001, F=33.43, df=495) when compared with one-way ANOVA. For further analysis, we selected the random forest regression model as it had the lowest median error (Figure 3a-c). Random forest models can also represent complex interactions among variables, which linear regressions cannot do, and may perform better in datasets where the number of features approaches or exceeds the number of observations (Breiman 2001). We term the outcome of this model FRIGHT age for <u>Fr</u>ailty Inferred <u>G</u>eriatric <u>H</u>ealth <u>T</u>imeline.

**Figure 3.**
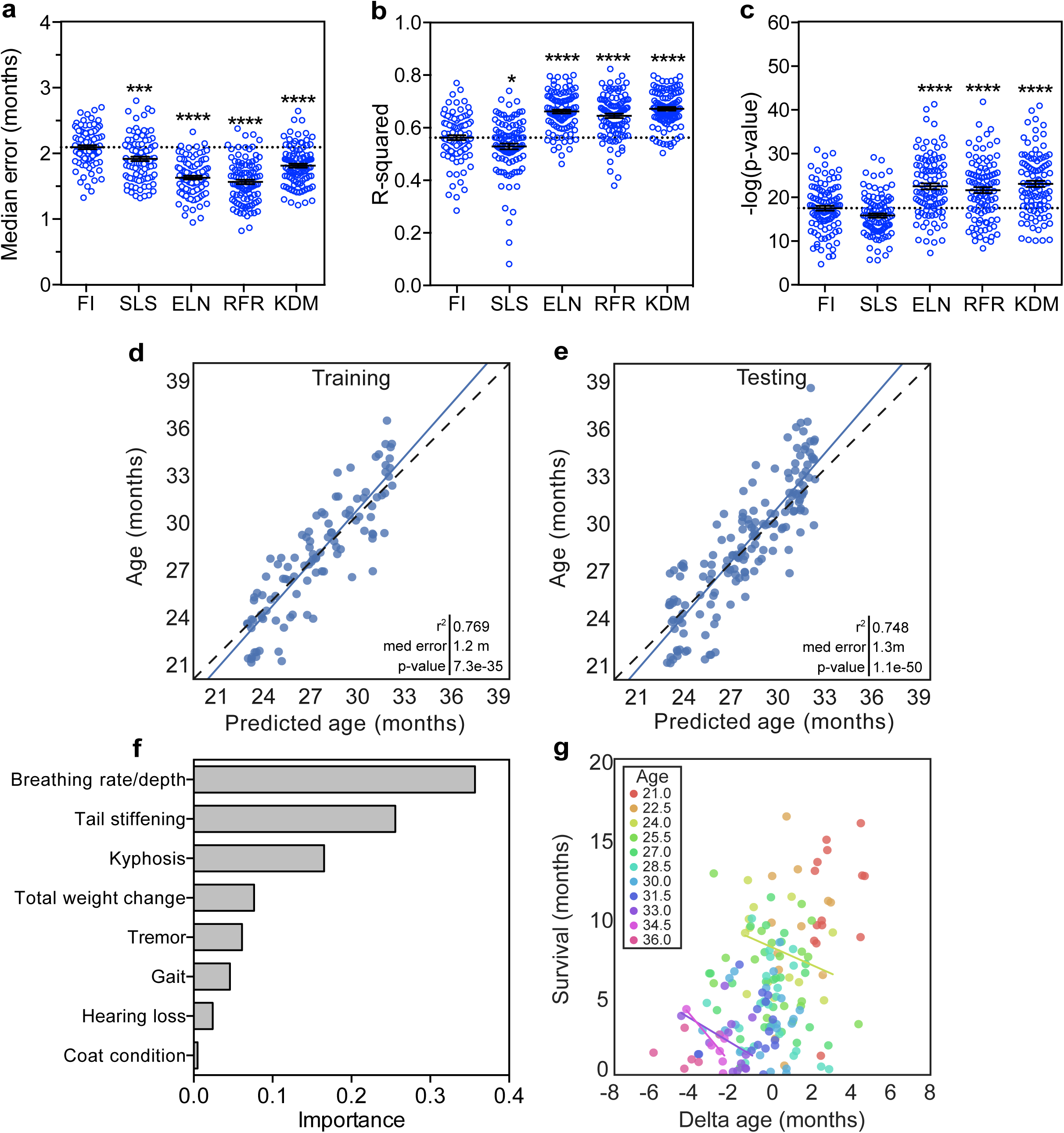
Multivariate regressions of individual FI items to predict age (FRIGHT age). (a-c) Median error, r-squared values and p-values for univariate regression of frailty index (FI) score, and multivariate regressions of the individual FI items using either simple least squares (SLS), elastic net (ELN), the Klemera-Doubal method (KDM), or random forest regression (RFR) for chronological age in the mouse training set. All models were tested with bootstrapping with replacement repeated 100 times, and each bootstrapping incidence is plotted as a separate point. **** indicates p-value <0.0001 and *** indicates p-value <0.001 compared to FI model with one-way ANOVA. (d-e) Random forest regression of the individual frailty index items for chronological age on training and testing datasets. This model is termed FRIGHT (<u>Fr</u>ailty <u>I</u>nferred <u>G</u>eriatric <u>H</u>ealth <u>T</u>imeline) age. (f) Importances of top items included in the FRIGHT age model. (g) Residuals of the regression (delta age) plotted against survival for individual ages. Regression lines are only graphed for ages where there is an r^2^ value > 0.1.

When assessed on the testing dataset, FRIGHT age had a strong correlation with chronological age, with a median error of 1.3 months and an r-squared value of 0.748 (p=1.1e^-50^) (Figure 3d-e). The items that were the largest contributors to FRIGHT age included breathing rate, tail stiffening, kyphosis, and total weight change (Figure 3f). While FRIGHT age was superior to the FI score at predicting chronological age (Figure 3a-c), the error from the predictions (delta age) were not well-correlated with mortality (Figure 3g). For the majority of individual age groups the r^2^ values of the correlation between FRIGHT age and survival were < 0.1, indicating poor correlation (Table 1). Interestingly, the correlations were stronger for mice aged 34 months or greater, indicating that perhaps FRIGHT age is predictive of mortality only in the oldest mice (Table 1). This may be because the individual parameters that correlate well with chronological age are not necessarily the same as those that correlate well with mortality at all ages. Thus FRIGHT age has value as a predictor of apparent chronological age (eg. “this mouse looks thirty months old”) but it is not yet clear whether it can serve as a predictor of other age-related outcomes.

### Multivariate regressions of individual frailty items to predict life expectancy (AFRAID clock)

As FRIGHT age was not predictive of mortality at most ages, we sought to build a model based on individual FI items to better predict life expectency. We began by calculating the correlation between each individual parameter and survival (number of days from date of FI assessment to date of death). Chronological age was the best predictor of mortality (r^2^=0.35, p=1.9e^-27^), followed by FI score, tremor, body condition score, and gait disorders (Table 3). However, many of these individual parameters appear to be better predictors than they are, as a result of their covariance with chronological age. Their correlation with survival is largely only for mice of different ages, and not of the same age. Indeed, when these correlations were calculated for only mice of the same age (n=6-27 per individual age), the only parameters that reached significance at most ages were those measuring weight change (data not shown).

**Table 3.** Correlation coefficients (r^2^) and p values for individual frailty items with life expectancy.

| **Item** | **$r^2$** | **p value** |
| --- | --- | --- |
| Age (days) | 0.35 | <0.001 |
| FI Score | 0.31 | <0.001 |
| Tremor | 0.25 | <0.001 |
| Body condition score | 0.20 | <0.001 |
| Gait disorders | 0.19 | <0.001 |
| Tail stiffening | 0.19 | <0.001 |
| Breathing rate/depth | 0.17 | <0.001 |
| Hearing loss | 0.17 | <0.001 |
| Kyphosis | 0.12 | <0.001 |
| Distended abdomen | 0.11 | <0.001 |
| Menace reflex | 0.08 | <0.001 |
| % twc | 0.07 | <0.001 |
| Forelimb grip strength | 0.07 | <0.001 |
| Alopecia | 0.05 | <0.001 |
| Threshold % rwc | 0.04 | <0.001 |
| Microphthalmia | 0.03 | <0.001 |
| Body weight score | 0.03 | <0.001 |
| Coat condition | 0.03 | <0.001 |
| Diarrhea | 0.03 | 0.01 |
| Mouse grimace scale | 0.02 | 0.01 |
| Loss of fur color | 0.02 | 0.01 |
| Dermatitis | 0.02 | 0.03 |
| Piloerection | 0.02 | 0.03 |
| % rwc | 0.01 | 0.06 |
| Rectal prolapse | 0.01 | 0.07 |
| Penile prolapse | 0.01 | 0.11 |
| Tumors | 0.01 | 0.13 |
| Eye discharge swelling | 0.01 | 0.17 |
| Vestibular disturbance | 0.01 | 0.19 |
| Vision loss | 0.01 | 0.27 |
| Cataracts | 0.00 | 0.38 |
| Corneal capacity | 0.00 | 0.44 |
| Loss of whiskers | 0.00 | 0.51 |
| Nasal discharge | 0.00 | 1.00 |

Twc, total weight change; rwc, recent weight change.

To build a model to predict mortality, we trained a regression using FI as a single variable, and multivariate regressions using the FI items and chronological age with the simple least squares, elastic net, and random forest methods. As before, we compared these models using bootstrapping on the training set, and one-way ANOVA with Dunnett’s posthoc test of r^2^ value, p-value and median error (Figure 4a-c). For prediction of survival, a random forest regression (RFR) model was the superior model with higher r^2^ value, lower p-value and lower median error than any of the other models (Figure 4a-e, Figure S4). We termed the outcome of this model the AFRAID clock for <u>A</u>nalysis of <u>Frai</u>lty and <u>D</u>eath. The most important variables in the model were total weight loss, chronological age, and tremor, followed by distended abdomen, recent weight loss, and menace reflex (Figure 4f). In the testing dataset, the AFRAID clock was well-correlated with survival (r^2^=0.505, median error = 1.7 months, p=1.1^-26^) (Figure 4e). The AFRAID clock was also correlated with survival at individual ages (Figure 4g) with r^2^ >0.1 and p value < 0.05 at 24, 27, 28.5, 30 and 34.5 months of age (Table 1). Plotting the survival curves of mice with the lowest and highest AFRAID clock scores at given ages, as determined by the top and bottom quartiles, demonstrated a clear association with mortality risk for all age groups (Figure 4h-k). These results suggest that the AFRAID clock may be useful for comparing the lifespan effects of interventional studies in mice many months before their death.

**Figure 4.**
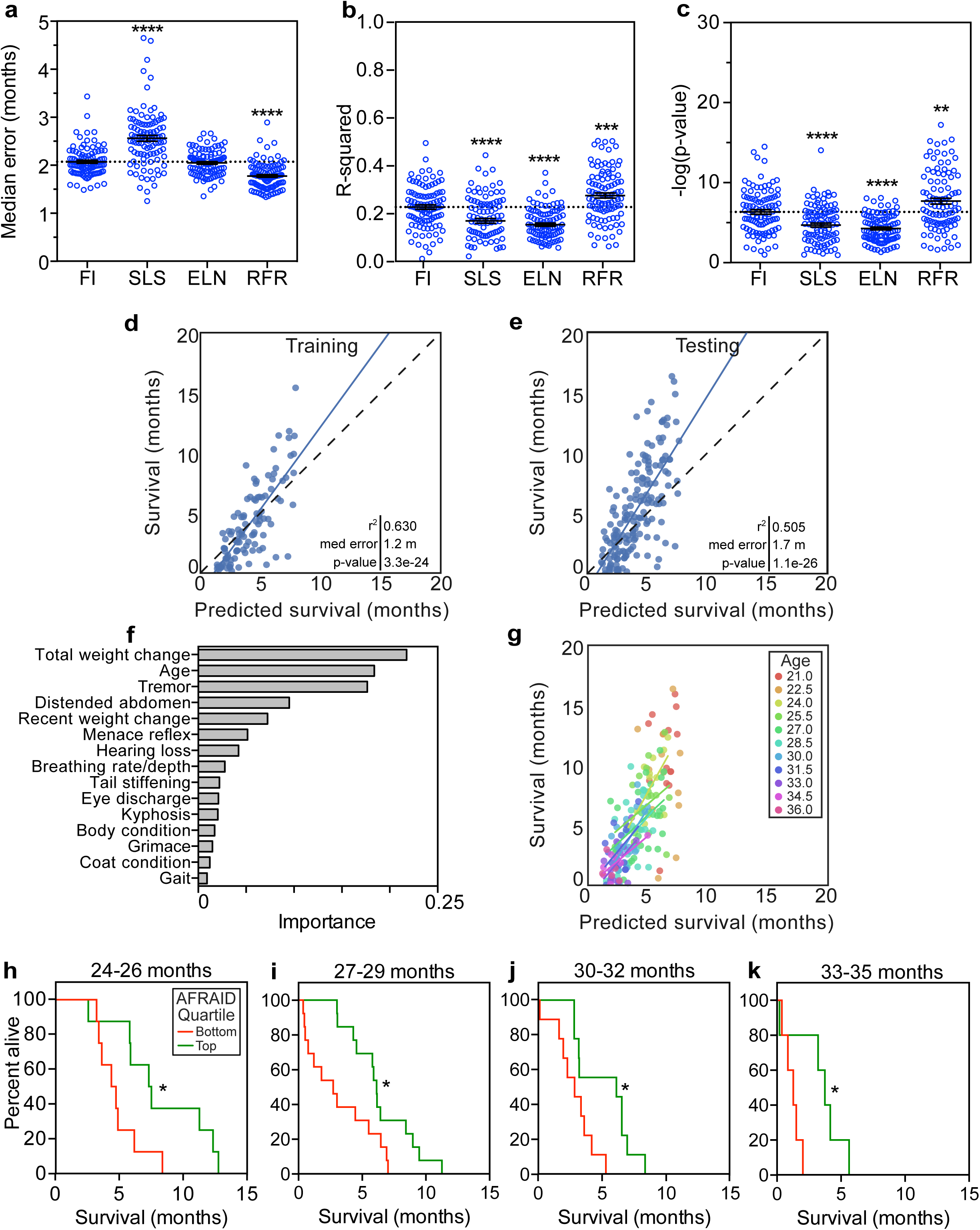
Multivariate regressions of individual FI items to predict life expectancy (AFRAID Clock). (a-c) Median error, r-squared values and p-values for univariate regression of frailty index (FI) score, and multivariate regressions of the individual FI items using either simple least squares (SLS), elastic net (ELN) or random forest regression (RFR) for life expectancy in the mouse training set. All models were tested with bootstrapping with replacement repeated 100 times, and each bootstrapping incidence is plotted as a separate point. **** indicates p-value <0.0001 and *** indicates p-value <0.001 compared to FI model with one-way ANOVA. (d-e) Random forest regression of the individual frailty index items for life expectancy on training and testing datasets, plotted against actual survival. This model is termed the AFRAID (<u>A</u>nalysis of <u>Frai</u>lty and <u>D</u>eath) clock. (f) Importances of top items included in the AFRAID clock. (g) AFRAID clock scores plotted against actual survival for individual mouse age groups in the testing dataset. Regression lines are only graphed for ages where there is an r^2^ value > 0.1. (h-k) Kaplan-Meier curves of the bottom and top quartiles of AFRAID clock scores for mice over 1-2 assessments at 24-26, 27-29, 30-32 and 33-35 months of age. * indicates p-value <0.05 compared with the log-rank test.

### Analysis of FRIGHT age and AFRAID clock in response to interventions

One ultimate utility for biological age models would be to serve as early biomarkers for the effects of interventional treatments, which are expected to extend or reduce healthspan and lifespan. A recently published study measured FI in 23-month old male C57BL/6 mice treated with the angiotensin converting enzyme (ACE) inhibitor enalapril (n=21) from 16 months of age, or age-matched controls (n=13) (Keller et al. 2018). As previously published, enalapril reduced the average FI score compared to control treated mice (Figure 5a). When FRIGHT age was calculated for these mice, the enalapril treated mice appeared to be a month younger than the control mice (control 27.8±1.1 months; enalapril 26.8±1.4 months, p=0.046, t=2.1, df=32)(Figure 5b). When the data were converted to a prediction of survival with the AFRAID clock, the enalapril treated mice were not predicted to live longer (control 5.9±0.7 months; enalapril 6.2±0.9 months, p=0.24, t=1.09, df=32) (Figure 5c). This is interesting in light of the fact that enalapril has been shown to improve health, but not maximum lifespan, in mice (Keller et al. 2018; Harrison et al. 2009).

**Figure 5.**
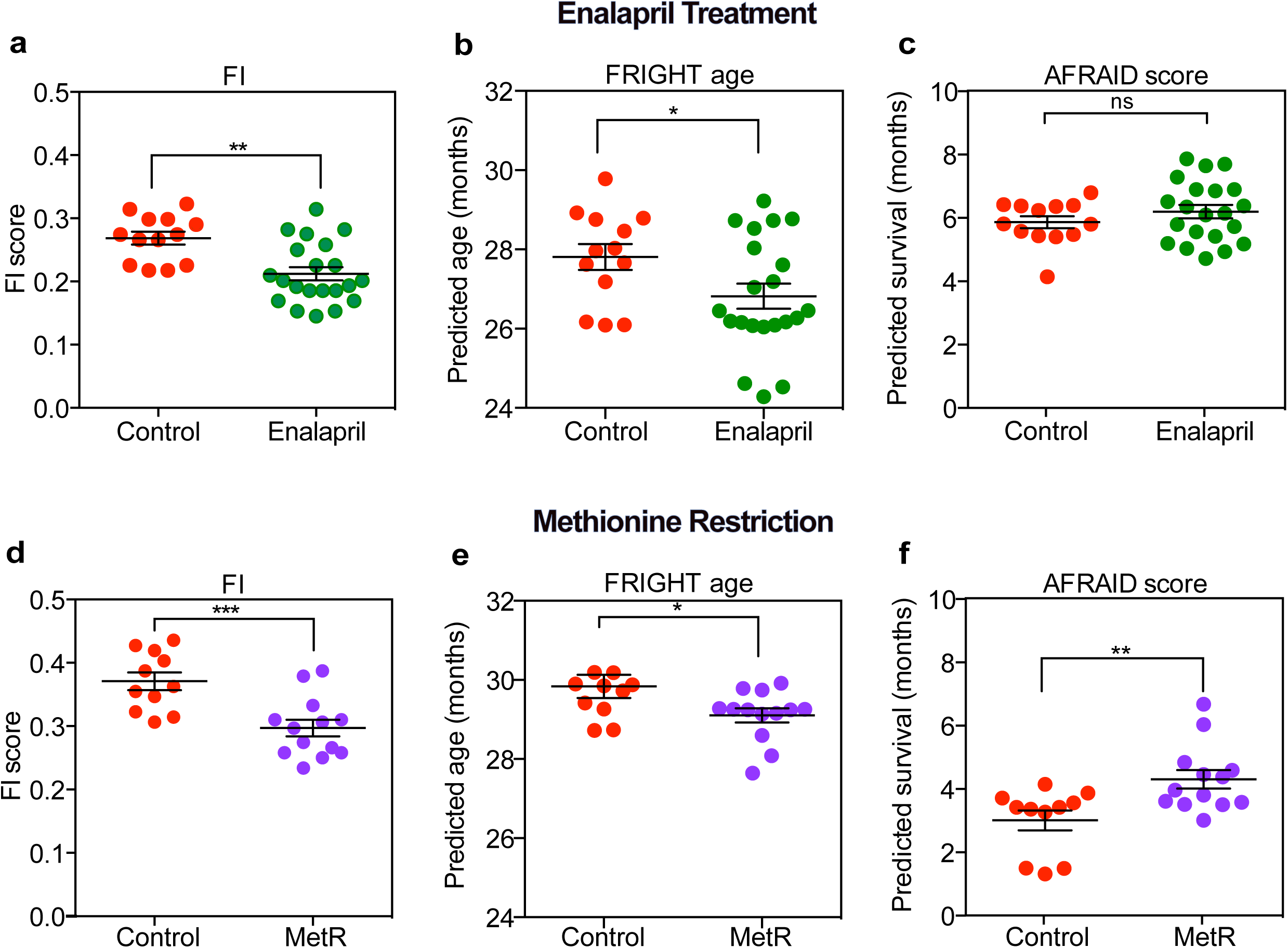
Response of FRIGHT age and AFRAID clock to interventions. (a-c) FI score, FRIGHT age and AFRAID clock for male 23 month old C57BL/6 mice treated with enalapril-containing food (280 mg/kg) or control diet from 16 months of age. Data reanalyzed from previously published work (Keller et al. 2018). (d-f) FI score, FRIGHT age and AFRAID clock for male 27 month old C57BL/6 mice treated with either a control diet (0.45% methionine) or methionine restricted diet (0.1% methionine) from 21 months of age. * indicates p-value <0.05, ** p-value less than <0.01, and *** p-values less than <0.001 compared with independent t-tests.

Methionine restriction is a robust intervention that extends the healthspan and lifespan of C57Bl/6 mice (Miller et al. 2005; Orentreich et al. 1993; Sun et al. 2009). We placed mice on a methionine restriction (0.1% methionine, n=13) or control (n=11) diet, from 21 months of age. We assessed frailty at 27 months of age and calculated FI, FRIGHT age and AFRAID clock. The methionine restricted mice had significantly lower FI scores (control 0.37±0.30; MR 0.30±0.04, p=0.0009, t=3.8, df=22)(Figure 5d), as well as a FRIGHT age 0.7 months younger than control-fed mice (control 29.8±0.9 months; MR 29.1±0.6 months, p=0.039, t=2.19, df=22)(Figure 5e). Using the AFRAID clock, the methionine restricted mice were predicted to live 1.3 months longer than controls (control 3.0±1.0 months; enalapril 4.3±1.0 months, p=0.006, t=3.02, df=22)(Figure 5f). These analyses demonstrate that the FRIGHT age and AFRAID clock models are responsive to healthspan and lifespan extending interventions.

## DISCUSSION

This is the first study to measure frailty longitudinally in a population of naturally aging mice that were tracked until their natural deaths in order to predict healthspan and lifespan. We show that the frailty index is not only correlated with but is also predictive of both age and survival in mice, and we have used components of the frailty index to generate two clocks: FRIGHT age, which models apparent chronological age better than the frailty index itself, and the AFRAID clock, which predicts life expectancy with greater accuracy than the frailty index. In essence, FRIGHT age is an estimation of how old a mouse appears to be, and the AFRAID clock is a prediction of how long a mouse has until it dies (a death clock). Finally, FRIGHT age and the AFRAID clock were shown to be sensitive to two healthspan or lifespan increasing interventions: enalapril treatment and dietary methionine restriction.

The major advantage of the frailty index, and our models of the frailty index items, as aging biometrics is their ease of use. FI is quick and essentially free to assess, requires no specialized equipment or training, and has no negative impact on the health of the animals. We encourage future longevity studies to incorporate periodic frailty assessments as a routine measure into their protocols. This will help further determine the utility of frailty itself, as well as our FRIGHT age and AFRAID clock models, for predicting outcomes of interest, and may eventually be used as a screening tool to decide whether to continue expensive interventional longevity studies after a short duration. Additionally, use of these non-invasive frailty measures in longevity studies will enable researchers to detect not only possible changes in lifespan, but also healthspan, arguably a more important outcome. We have made our clock calculators available online (https://github.com/SinclairLab/frailty).

The recently developed DNA methylation clocks are also promising biomarkers of biological age. In humans, these clocks are highly correlated with chronological age, and are able to predict, at the population level, mortality risk and risk of age-related diseases (Horvath and Levine 2015; Horvath et al. 2015, 2016; Horvath and Raj 2018; Quach et al. 2017; Maierhofer et al. 2017; Levine et al. 2018). Methylation clocks have also been developed for mice, and shown to correlate with chronological age, and respond to lifespan-increasing interventions such as calorie restriction (Meer et al. 2018; Petkovich et al. 2017; Cole et al. 2017; Stubbs et al. 2017), but their association with mortality has not yet been explored. However, the major drawback of these mouse clocks is that they require repeated invasive blood collections and time-consuming and expensive data acquisition and analysis procedures.

This is the first time, to our knowledge, that frailty has been used to predict individual life expectancy in either humans or mice. Previous predictive mortality measures in mice have either focused on the acute prediction of death such as in the context of sepsis (Trammell and Toth 2011; Ray et al. 2010), focused on only a few measures resulting in low or moderate correlations with survival (Ingram et al. 1982; Miller 2001; Miller et al. 2002; Harper et al. 2003; Fahlström, Zeberg, and Ulfhake 2012; Swindell, Harper, and Miller 2008), or used short-lived mouse strains (Martínez de Toda et al. 2019). The AFRAID clock, which was modelled in the commonly used C57BL/6 mouse strain and includes 33 variables, is able to predict mortality with a median error of 46 days across multiple ages. The real value of a biological age measure for mice, however, is in predicting how long individual mice of the same chronological age will live. The AFRAID clock was also able to predict mortality at specific ages, even as early as 24 months (approximately six months before the average lifespan, and twelve months before maximum lifespan without intervention). This provides exciting evidence that this measure could be used in interventional longevity studies to understand whether an intervention is working to delay aging at an earlier timepoint than death.

Indeed, we show in the current study that treatment with the angiotensin-converting enzyme (ACE) inhibitor enalapril reduced FRIGHT age compared to controls but did not change the AFRAID clock. Enalapril is known to increase healthspan but not lifespan (Harrison et al. 2009), indicating the value of these measures in detecting healthspan improvements even in the absence of an increase in lifespan. The dietary intervention of methionine restriction is known to increase healthspan and lifespan (Miller et al. 2005; Orentreich et al. 1993; Sun et al. 2009), and we saw reduced FRIGHT age and increased AFRAID clock scores in methionine restricted mice at 27 months compared to controls. This means that had this been a longevity study, these measures would have given an indication of the lifespan outcomes less than halfway through the predicted study timeframe.

Studies in humans have used the frailty index to determine increased risk of mortality within specific time periods (Song, Mitnitski, and Rockwood 2010; Kane et al. 2017; Blodgett et al. 2015), but not to predict individual life expectancies. In theory the AFRAID clock could be easily adapted to predict mortality from human frailty index data. This has likely not been done as of yet, as it would require a large dataset that includes longitudinal assessments of frailty index items with mortality follow-up. This type of study is rare, particularly in an aging population. Even large cohort studies such as NHANES do not include enough people aged over 80 to allow for their specific ages to be released due to risk of identification. It would be interesting in future research to apply machine learning algorithms such as those used in the current study to predict individual life expectancy using frailty index data in humans.

The aim of all three frailty metrics presented here, FI score, FRIGHT age, and the AFRAID clock, are robust methods for the appraisal of biological age. Without a clear biomarker with which to compare these three metrics, an assessment of their relative value is difficult. In one sense, FRIGHT age is the best because it tracks most closely with chronological age, in spite of its lack of sensitivity to predict mortality. An intervention that slows aging would likely suppress all aspects of aging including those that don’t impact life expectancy (for example hair greying) and FRIGHT age would detect such changes. In another sense, the AFRAID clock is the superior metric, because an increase in life expectancy, median and maximum, is the current benchmark for the success of an aging intervention. One could also argue that overall unweighted frailty index is the best metric. While it is not best at predicting either chronological age or mortality, it is better than either FRIGHT age or AFRAID clock at predicting both. The best approach may be to employ all three estimates.

The predictive power of these models for both age and lifespan could be improved by the inclusion of larger n values, and the assessment of frailty from ages younger than 21 months. Additionally we could model biological age markers, not to predict chronological age or mortality alone, but rather a more complex composite measure of age-associated outcomes. Indeed, DNA methylation clocks that are trained on surrogate biomarker and biometrics for mortality including blood markers and plasma proteins plus gender and chronological age (Levine et al. 2018; Lu et al. 2019) seem to have greater predictive power than those modelled on chronological age or mortality alone (Horvath 2013a; Zhang et al. 2017). Future studies could develop a new model based on the frailty items assessed here but modelled to predict a composite outcome including physiological measures in addition to chronological age. The models discussed in this study could also benefit from the incorporation of additional input variables, especially from relatively non-invasive molecular and physiological biomarkers or biometrics. Much can be inferred from tallying gross physiological deficits as has been done here with the mouse frailty index. However, these deficits have cellular and molecular origins which may add predictive value at much earlier time points if they can be identified. Frailty indices based on deficits in laboratory measures such as blood tests can detect health deficits before they are clinically apparent in both humans and mice (Howlett et al. 2014; Kane et al. 2018). Still, even after the development of such composite clocks, the metrics described here – FI, FRIGHT age, and the AFRAID clock – will serve as rapid, non-invasive means to assess biological age and life expectency, accelerating and augmenting studies to identify interventions that improve healthspan and lifespan.

## METHODS

### Animals

All experiments were conducted according to protocols approved by the Institutional Animal Care and Use Committee (Harvard Medical School). Aged males C57BL/6Nia mice were ordered from the National Institute on Aging (NIA, Bethesda, MD). A cohort of mice (n=27) were injected with AAV vectors containing GFP as a control group for a separate longevity experiment at 21 months of age. This did not affect their frailty or longevity in comparison to the rest of the mice (n=24), which were untreated (Supplementary Figure 1). Both sets of animals had normal median (954 and 922 days) and 90^th^ percentile (1125 and 1073 days) lifespans, slightly surpassing those cited by Jackson Labs (median 878 days, maximum 1200 days) (Fasting 1979; Kunstyr and Leuenberger 1975), demonstrating that the mice were maintained and aged in healthy conditions. Mice were only euthanized if determined to be moribund by an experienced researcher or veterinarian based on exhibiting at least two of the following: inability to eat or drink, severe lethargy or persistent recumbence, severe balance or gait disturbance, rapid weight loss (>20%), an ulcerated or bleeding tumor, and dyspnea or cyanosis. In these rare cases, the date of euthanasia was taken as the best estimate of death.

### Mouse Frailty Assessment

Frailty was assessed longitudinally by the same researcher (A.K.), as modified from the original mouse clinical FI (Whitehead et al 2014, Kane et al., 2017). Malocclusions and body temperature were not assessed in the current study, so a frailty index of 29 total items was used. Individual frailty index parameters are listed in Supplementary Figure 1. Briefly, mice were scored either 0, 0.5 or 1 for the degree of deficit they showed in each of these items with 0 representing no deficit, 0.5 representing a mild deficit and 1 representing a severe deficit. For regression analyses, prediction variables were added to represent body weight change: total percent weight change, from 21 months of age; recent percent weight change, from one month before the assessment; and threshold recent weight change – mice received a score for this item if they gained more than eight grams or lost more than ten grams from the previous month. FI scoresheet for automated data entry (Figure S1g) is available online (https://github.com/SinclairLab/frailty).

### Intervention studies

#### Enalapril treatment

Data from enalapril treated mice were reanalyzed from previously published work (Keller et al. 2018). Briefly, male C57BL/6 mice purchased from Charles River mice were treated with control or enalapril food (30 mg/kg/day) from 16 months of age and assessed for the frailty index at 23 months of age.

#### Methionine Restriction

Male C57BL/6Nia mice were obtained from the NIA at 19 months of age and fed either a control diet (0.45% methionine) or methionine restricted diet (0.1% methionine) from 21 months of age. Custom mouse diets were formulated at research diets (New Brunswick, NJ) (catalog #’s A17101101 and A19022001). Mice were assessed for the frailty index at 27 months of age.

### Modelling and Statistics

All analysis was done in Python. Training and testing datasets were randomly split 50:50 and were separated by mouse rather than by assessment. Missing frailty data (18 individual data points out of 2460 total data points) was replaced by the median value for that item for that age group. All models were assessed with bootstrapping with replacement, repeated 100 times. The fit of the models was determined with the r^2^ value which determines the proportion of the variance in our predicted outcome that is explained by the model, the median residual/error which represented the median difference between the actual and predicted outcome values, and the p value of the regressions. Median error, r-squared and p values were compared across measures of FRIGHT age or AFRAID clock (Figure 3A-C and Figure 4A-C) with one-way ANOVA and Dunnett’s posthoc test. Kaplan-Meier survival curves of the highest and lowest quartiles of AFRAID clock scores (Figure 4) were compared with the log-rank test. FI, FRIGHT age and AFRAID clock scores across intervention and control groups (Figure 5) were compared with independent samples t-tests. For all statistics, p values less than 0.05 were considered significant. All data is presented as mean ± SD, except error bars on figures indicate standard error of the mean.

#### Least Squared and Elastic Net Regressions

Regressions were performed using algorithms provided in the Scikit-learn package (Pedregosa et al. 2011) in Python. Least squared regression was performed using the standard LinearRegression algorithm. Elastic net was performed with the ElasticNet algorithm with coefficients restrained as positive.

#### Klemera-Doubal Biological Age Method

We calculated Klemera-Doubal biological age of each mouse using the methods first described by Klemera and Doubal (Klemera Doubal 2006) and later demonstrated by Levine (Levine 2013) and Belsky (Belsky 2015). The Klemera-Doubal Method (KDM) uses multiple linear regression but improves upon this by reducing multicollinearity between biological variables, which are intrinsically correlated. The KDM method consists of *m* regressions of age against each of *m* predictors. A basic biological age is then predicted based on the following equation:

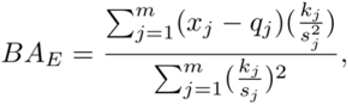

where *k*_*j*_, *q*_*j*_, and *s*_*j*_ represent the slope, intercept, and root mean square error of each of the *m* regressions, respectively. While Klemera and Doubal further suggest using chronological age as a corrective term to limit the bounds of each predicted value, we used the version of the algorithm without age as (1) predictions of mortality on an ROC curve using both the corrected and uncorrected biological age showed little difference (data not shown), and (2) for the purposes of this study, we wanted to demonstrate the utility of the variables alone as predictors of age without knowledge of the true chronological age of the mouse.

#### Random Forest

Random forests are a type of machine learning algorithm which combines many decision trees into one regression outcome (Breiman 2001). Random forest modelling was performed using the Scikit-learn RandomForestRegressor algorithm (Pedregosa et al. 2011). Models were made with 1000 trees, and the minimum number of samples required for a branch split was limited to prevent overfitting. We also computed and plotted the feature importance for each of the items with the highest value for this outcome. Feature importance is the amount the error of the model increases when this item is excluded from the model.

## ACKNOWLEDGMENTS

This work was supported by the Glenn Foundation for Medical Research and grants from the NIH (R37 AG028730, R01 AG019719, R01 DK100263, R01 DK090629-08), and Epigenetics Seed Grant (601139_2018) from Department of Genetics, Harvard Medical School. A.E.K is supported by an NHMRC CJ Martin biomedical fellowship (GNT1122542). Grants to SEH from the Canadian Institutes for Health Research (PGT 162462) and the Heart and Stroke Foundation of Canada (G-19-0026260). Grant to JM from the NIH (2R56AG036712-06A1). D.A.S. is a founder, equity owner, advisor to, director of, consultant to, investor in and/or inventor on patents licensed to Vium, Jupiter Orphan Therapeutics, Cohbar, Galilei Biosciences, GlaxoSmithKline, OvaScience, EMD Millipore, Wellomics, Inside Tracker, Caudalie, Bayer Crop Science, Longwood Fund, Zymo Research, Immetas, and EdenRoc Sciences (and affiliates Arc-Bio, Dovetail Genomics, Claret Bioscience, Revere Biosensors, UpRNA and MetroBiotech, Liberty Biosecurity). Life Biosciences (and affiliates Selphagy, Senolytic Therapeutics, Spotlight Biosciences, Animal Biosciences, Iduna, Continuum Biosciences, Jumpstart Fertility (an NAD booster company), and Lua Communications). Iduna is a cellular reprogramming company, partially owned by Life Biosciences. DS sits on the board of directors of both companies. D.A.S. is an inventor on a patent application filed by Mayo Clinic and Harvard Medical School that has been licensed to Elysium Health; his personal share is directed to the Sinclair lab. For more information see https://genetics.med.harvard.edu/sinclair-test/people/sinclair-other.php. MSB is a stockholder for MetroBiotech and Animal Biosciences, a division of Lifebiosciences. Other authors have no conflicts to declare.

## SUPPLEMENTARY FIGURE LEGENDS

**Figure S1.**
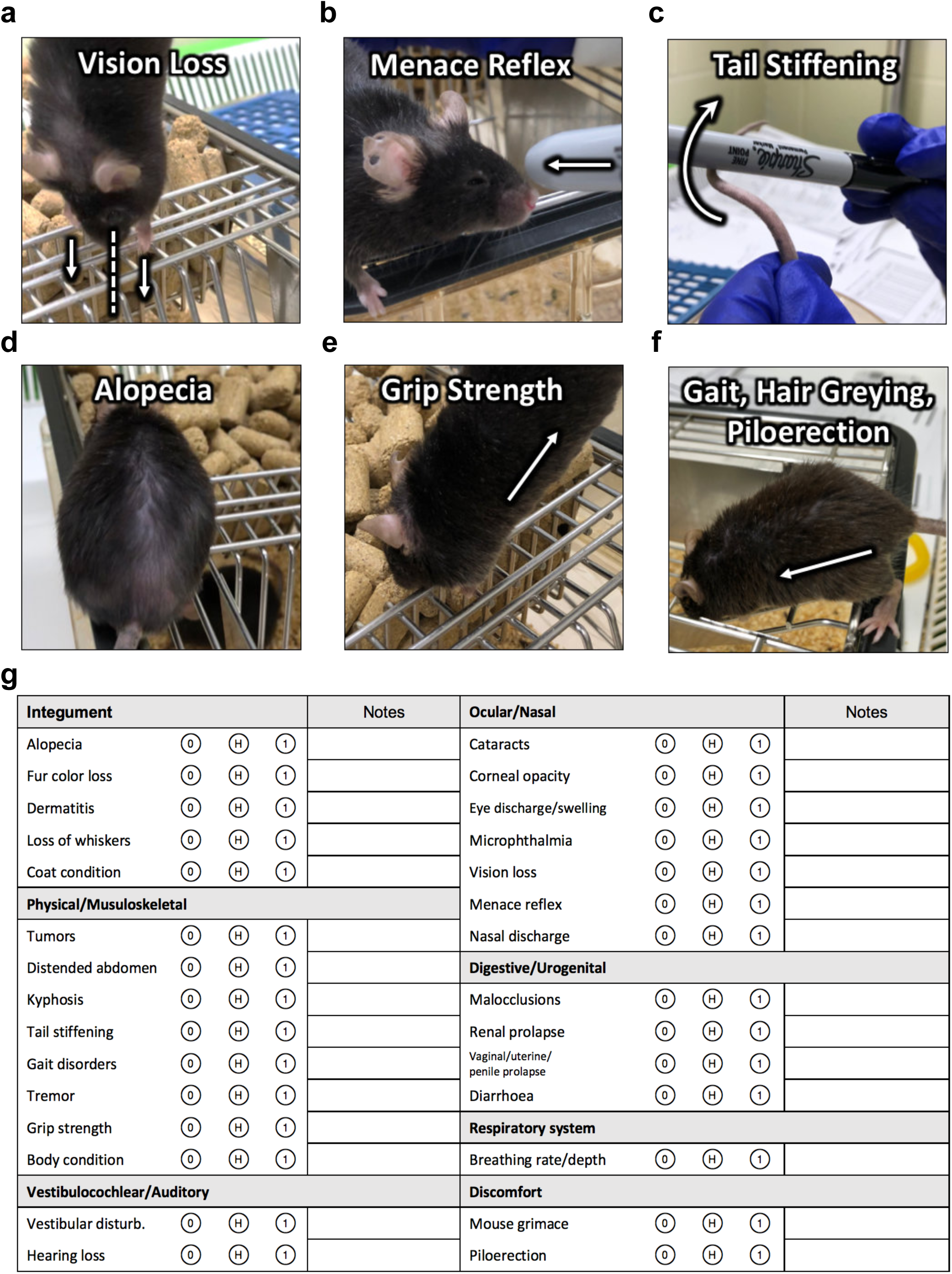
Frailty Index (FI) items. (a-f) Examples of deficits measured with the non-invasive mouse clinical FI assessment (Whitehead et al. 2014). (g) Scoresheet for automated data entry of FI item scoring, modified from original paper (available at https://github.com/SinclairLab/frailty).

**Figure S2.**
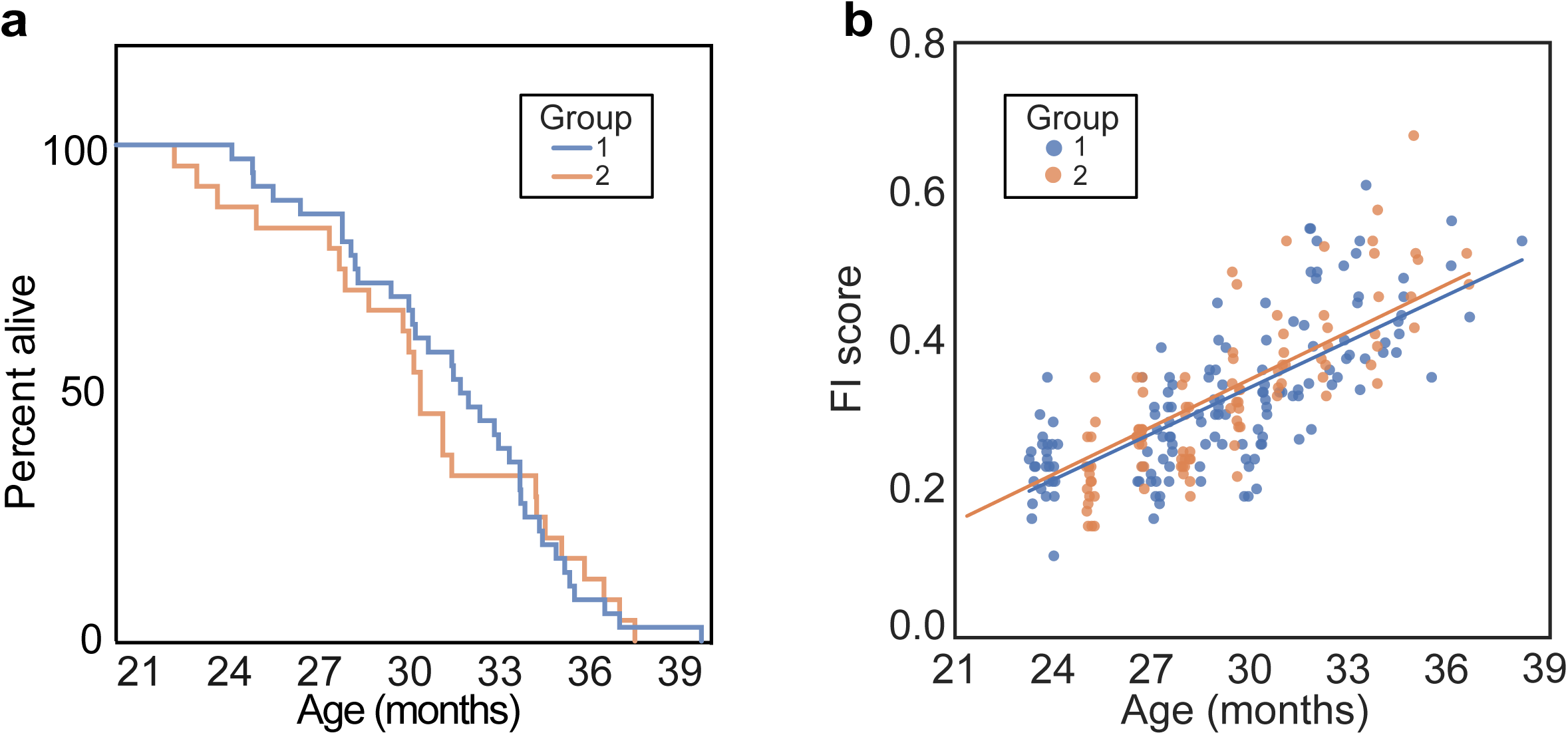
Mortality curves and frailty index for mice cohorts. (a) Kaplen-Meier curves for male C57BL/6 mice included in the current study. One cohort of mice (blue, n=27) was injected with AAV vectors containing GFP as a control group for a separate longevity experiment, while the second cohort (orange, n=24) was untreated. (b) All FI scores from 21 months of age for mice included in the current study.

**Figure S3.**
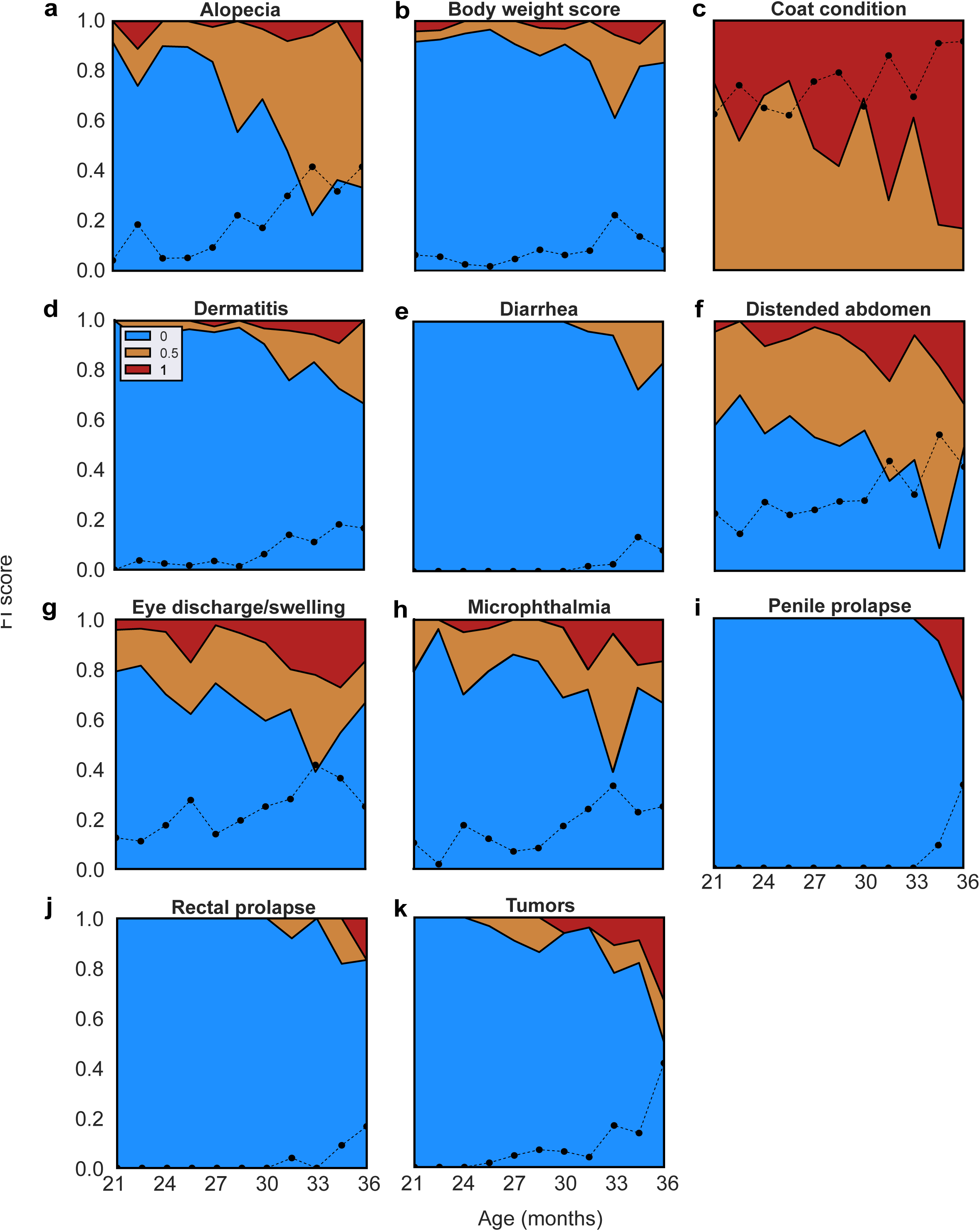
Individual FI items vary in their correlation with age. Mean scores across all mice (black line) for individual items of the frailty index from 21 to 36 months of age that had any positive correclation with age. Colors indicate proportion of mice at each age with each score (0, blue; 0.5, orange, 1, red).

**Figure S4.**
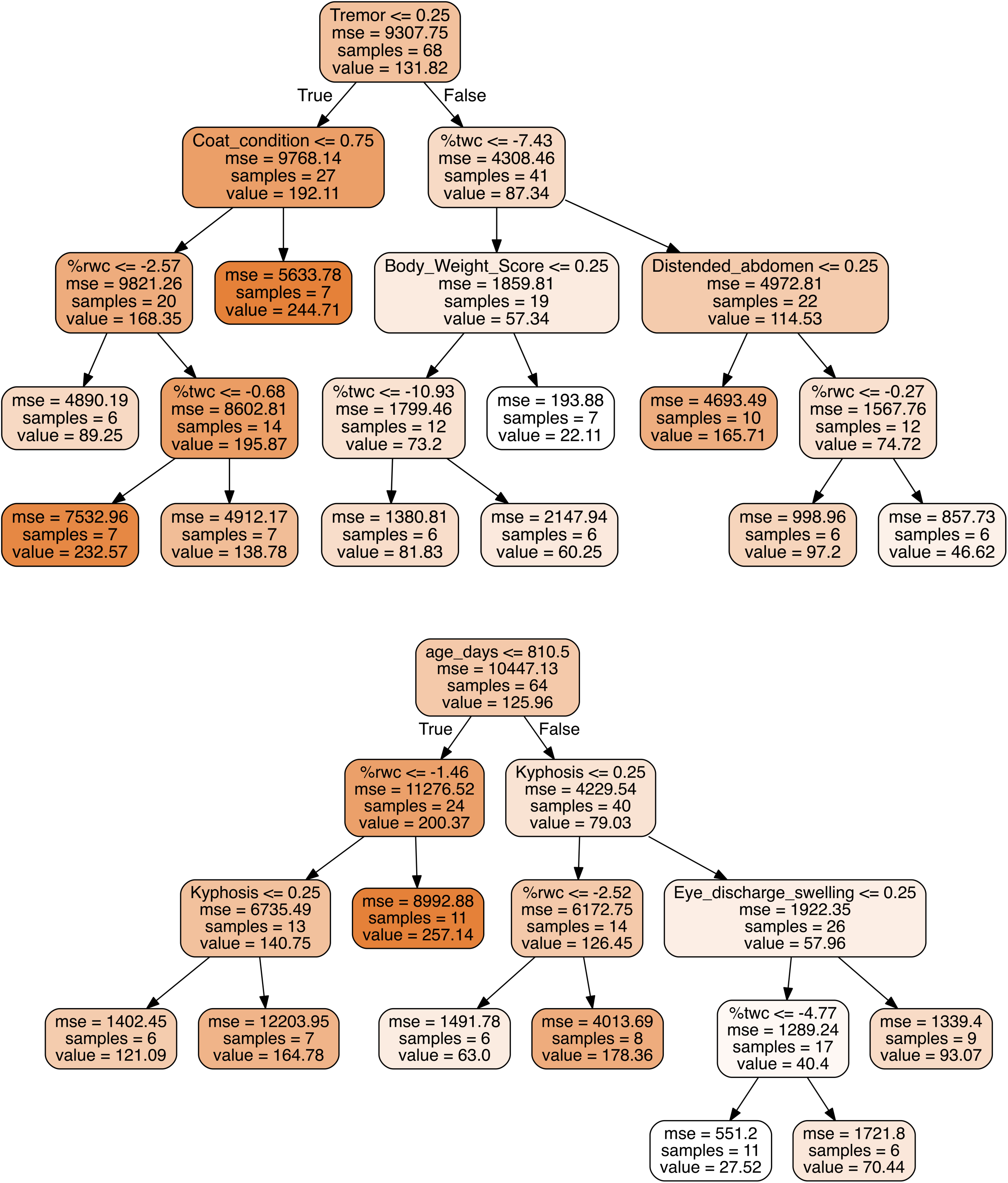
Example tree from random forest analysis. An illustration of two decision tree, out of the one thousand that comprise the AFRAID clock measure.

